# Current forest carbon offset buffer pools do not adequately insure against disturbance-driven carbon losses

**DOI:** 10.1101/2024.03.28.587000

**Authors:** William R. L. Anderegg, Anna T. Trugman, German Vargas, Chao Wu, Linqing Yang

## Abstract

Nature-based climate solutions in Earth’s forests could strengthen the land carbon sink and contribute to climate mitigation, but must adequately account for climate risks to the durability of carbon storage. Forest carbon offset protocols use a ‘buffer pool’ to insure against disturbance risks that may compromise durability. However, current buffer pool tools and allocations are not based on existing scientific data or models. Here, we use a tropical forest stand biomass model and an extensive set of long-term tropical forest plots to test whether current buffer pools are adequate to insure against observed disturbance regimes. We find that forest age and disturbance regime both influence necessary buffer pool sizes. In the vast majority of disturbance scenarios, current buffer pools are substantially smaller than required by carbon cycle science. Buffer pool estimates urgently need to be updated based on rigorous, open scientific datasets for nature-based climate solutions to succeed.

**Plain Language Summary:** Forests could contribute to climate mitigation through conservation and restoration activities. Carbon offsets are a widespread pathway to fund these nature-based climate solutions in forests, but must account for the risks to durability that forests face in a changing climate. Current carbon offset protocols have a buffer pool to insure against risk in different disturbance regimes, but the buffer pool contributions have not been tested with observed disturbance regimes and rigorous models. We tested these contributions using widespread tropical forest plot data and a carbon cycle model and find that the current buffer pool contributions are generally not adequate for most disturbance regimes. Our results highlight that better datasets, models, and tools are urgently needed in forest carbon offset protocols.

**Key points:**

- Nature-based climate solutions in forests face substantial and rising climate risks to durability
- Carbon offsets use a buffer pool to insure against disturbance, which is not currently based on rigorous evidence
- Our results reveal current carbon offset protocols do not have an adequate buffer pool for most tropical forest disturbance regimes

## 1. Introduction

Earth’s forests currently serve as a substantial carbon sink, absorbing roughly a quarter of human carbon emissions annually from the atmosphere (Bonan, 2008; Pan et al., 2011; Pugh et al., 2019). Alongside critical and necessary efforts to dramatically reduce fossil fuel emissions, forests can contribute to climate change mitigation as ‘nature-based climate solutions’ (NbCS), which are a suite of potential changes in management decisions to increase forest carbon stocks (Griscom et al., 2017; Nolan et al., 2021; Seddon, 2022). The most common forest-based NbCS to date are i) avoided conversion/loss where forests are protected from degradation or deforestation, ii) improved forest management where management changes are implemented to increase carbon stocks, and iii) afforestation/reforestation where forests are actively planted and restored (Griscom et al., 2017). The potential for NbCS is considered to be largest in tropical forests where there is widespread interest in leveraging NbCS efforts to reduce and stop deforestation through Reducing Emissions from Deforestation and Degradation (REDD+) efforts (Griscom et al., 2020; Haya, Alford-Jones, et al., 2023; Roopsind et al., 2019). Despite the potential for NbCS, widespread quality concerns about ongoing efforts (W. R. Anderegg et al., 2020; Badgley, Chay, et al., 2022; Badgley, Freeman, et al., 2022; Coffield et al., 2022; Haya, Alford-Jones, et al., 2023; Stapp et al., 2023; West et al., 2023) highlight the urgent need of rigorous carbon cycle science underpinning the tools and protocols that fund and implement NbCS projects.

To serve effectively as climate mitigation, forest NbCS projects must achieve a level of durability in carbon storage and account for any potential carbon losses, frequently through a buffer pool approach. Individual projects contribute a portion of credits into a common buffer pool that serves as an insurance policy and then those credits are retired when a reversal (loss) of carbon occurs in a project. Durability is particularly crucial when the NbCS effort is funded as part of a carbon offset, because the carbon stored in the forest is intended to offset fossil fuel carbon emissions that warm the climate for centuries to millennia (Archer et al., 2009; Solomon et al., 2009) and must be generally sustained through when global temperatures peak to avoid excess warming (Cullenward et al., 2023; Matthews et al., 2022). A frequent timescale used as a gold standard for durability by NbCS project protocols is a 100-year time horizon (W. R. Anderegg et al., 2020; Haya, Evans, et al., 2023). For buffer pools to effectively insure project durability, they urgently need to accurately account for the frequency and severity of climate-sensitive disturbances such as fire or drought, and also the carbon cycle consequences of those disturbances.

Carbon offset protocols developed by carbon registries are the most prominent and widely used mechanism currently to fund forest NbCS projects (Haya, Evans, et al., 2023; Ruseva et al., 2017). The carbon registry Verra, the largest registry in the world in the voluntary carbon market, recently released a new tool for calculating buffer pool based on different disturbance severity and frequencies which is expected to have large influence on carbon markets and climate policy broadly. Previous Verra protocols have been criticized for buffer pools that were much smaller than currently available science (Haya, Alford-Jones, et al., 2023). Thus, rapid and urgent evaluation of the robustness of the new Verra tool is a key scientific need with enormous policy relevance.

Here, we use a forest carbon model parameterized with extensive data from 125 tropical forest plots across multiple countries to ask: 1) Do existing buffer pool estimates conform with predictions from tropical forest carbon cycle dynamics? 2) What buffer pool contributions would be needed to adequately insure different disturbance frequencies and severities, while accounting for a wide range in observed forest growth and mortality rates? 3) How does forest age affect buffer pool contributions and thus how might buffer pool contributions vary between reforestation (i.e. younger forests) or other project types (typically older forests)?

## 2. Methods

We simulated the carbon cycle dynamics of tropical forests in response to different disturbance scenarios articulated by Verra’s Non-Permanence Risk Tool version 4.2 (Table S1 reproduced from Table 10 in AFOLU Non-Permanence Risk Tool V4.2 (Verra, 2023)). This tool requires project developers to estimate the frequency (return time) and severity (carbon lost) of key natural disturbances including fire, pest and disease outbreaks, extreme weather events such as droughts and hurricanes, and geological risks such as earthquakes and volcanoes, and provides 25 categories of buffer pool contributions for each combination of frequency and severity for each relevant disturbance. The stated goal of a carbon offset buffer pool’s ‘natural risks’ contributions is to insure against carbon losses from disturbances, such that a portfolio of projects remains able to deliver the promised climate mitigation benefits. In other words, the protocol will likely fail from a durability point of view if biomass falls below the initial biomass plus the buffer pool levels across many projects, indicating that climate risks (termed ‘natural risks’ in the protocol) exceeded the allotted buffer pool. Note that offset protocols also must meet rigorous criteria in additionality, leakage, net climate impacts, and monitoring/verification to succeed for climate mitigation (Cook-Patton et al., 2023; Haya, Alford-Jones, et al., 2023; West et al., 2023). Currently, the buffer pool contribution numbers per disturbance category provided in Table 10 of the Non-Permanence Risk Tool (Table S1) are not based on scientific data to our knowledge (i.e. no citation presented in reference material).

The forest biomass model used here was developed at Barro Colorado Island in Panama to quantify tropical forest age-biomass dynamics across spatial scales using a range of growth and mortality rates (Knapp et al., 2022). This model is a statistical model that simulates distributions of biomass trajectories based on input rates of carbon gains (growth and recruitment) and carbon losses (mortality) at a range of spatial scales, whereby the change in aboveground biomass is described as a differential equation involving a biomass gain parameter and a mortality parameter (J. I. Fisher et al., 2008; Knapp et al., 2022). The model successfully recreates biomass distributions across spatial scales using the long-term plot data in Panama (Knapp et al., 2022) and is ideally suited for an examination of the impacts of age, growth rates, and disturbance frequency and severity on biomass and carbon trajectories in tropical forests. We used the published tropical forest carbon gain and carbon losses from mortality that comprises 125 long-term monitoring plots in tropical forests across North and South America (Yu et al., 2019). This allows us to quantify the estimated buffer pool contributions needed across a wide range of observed climate, soils, topography, and species composition.

We extracted the geographic location and project type (improved forest management) of the 256 currently active forest offset projects in Verra’s registry. We overlaid these locations on a global forest age map in 2010 at 1000m resolution (Besnard et al., 2021) and extracted an estimated mean forest age for each project based on the boundaries of project shapefiles or the 3*3 pixels with the centroid coordinates of the project as the center pixel when the project shapefile is not available. We note that these projects were developed under previous methodologies with different (and generally lower) buffer pool contribution rules (Haya, Alford-Jones, et al., 2023), but they provide a useful starting point for understanding the age distributions of different project categories for informing the model initiation. We plot current projects on a previously-generated global map of stand-replacing disturbance risk (W. R. L. Anderegg et al., 2022) to contextualize the range of integrated 100-year disturbance risk and return interval of stand-clearing disturbance.

We then used a Monte Carlo approach where we sampled across all carbon gain and carbon loss measurements of the 125 long-term tropical forest plots (925 plot-by-census combinations) with 1000 iterations per plot-by-census combination. In each Monte Carlo iteration, we sampled a uniform distribution with a given range of disturbance probabilities derived directly from Verra’s Table 10 scenarios and a uniform distribution that covered the range of carbon losses from disturbance from the same table of scenarios. This provides a probabilistic set of assessments that captures the wide range of potential biomass trajectories in a given disturbance regime over 100 years for a given tropical forest set of demographic rates (carbon gains and carbon losses) informed by observations. We first chose two representative ages for these Monte Carlo analyses: 1) forest age of 71 years, which was the mean age for reforestation/afforestation projects in Verra’s existing projects, and 2) forest age of 189 years old, the mean age for avoided conversion and improved forest management existing projects. Finally, to quantify the effects of age directly, we ran a set of scenarios with stand ages of 50, 100, 150, and 200 years old.

We plotted the biomass remaining (%) 100 years after project initiation, given its frequent policy relevance, and plotted variability across the stand biomass model Monte Carlo as boxplots, adding in lines for the mean and 20^th^ percentile biomass remaining, which would provide more conservative guidance for what a buffer pool contribution would need to successfully insure 80% of projects. Finally, Verra allows increases of buffer pool contributions due to climate risks by up to 40% based on IPCC ‘climate impact drivers’ (Ruane et al., 2022) that attempt to coarsely capture climate-driven amplification of durability risks. Verra also allows reductions of buffer pool contributions by up to 75% based on projects’ risk mitigation activities (Verra, 2023). Thus, we plot the full potential buffer pool contribution range in our figures as well. For example, if a project determined the baseline risk was a 5% buffer pool contribution for a given risk, the full range of potential buffer pool contributions could be 1.25-7%, based on justifications provided by the project developer.

## 3. Results

Current Verra forest carbon offset projects are located across all biomes, though heavily concentrated in tropical forests, and are exposed to a substantial level of risk for stand-replacing disturbances based on a previous-generated global dataset (Fig. 1). Stand biomass model inputs spanned a large range of carbon gains and carbon loss estimates in the 925 plot-by-census combinations in our tropical forest dataset (Fig. 1). After accounting for this large variation in observed growth and mortality rates in our model simulations, we found that severe carbon losses from disturbance have major impacts on the forest carbon storage in a given tropical forest after 100 years (e.g. Fig. 2). Here we provided an example for the simulated biomass trajectories from the Monte Carlo at a single forest site (i.e. combination of age, carbon gain, and carbon loss) over 100 years in the scenario with biomass losses of 50-70% from disturbance occurring once every 25-50 years. In this case, our stand model estimates that an average of 57% of initial biomass remains after 100 years. In contrast, the Verra buffer pool contribution for this scenario assumes that >96% of biomass is remaining on average. Above 93% of biomass scenarios from our model fall below the insured level in Verra’s buffer pool (blue line; Fig. 2).

**Figure 1:**
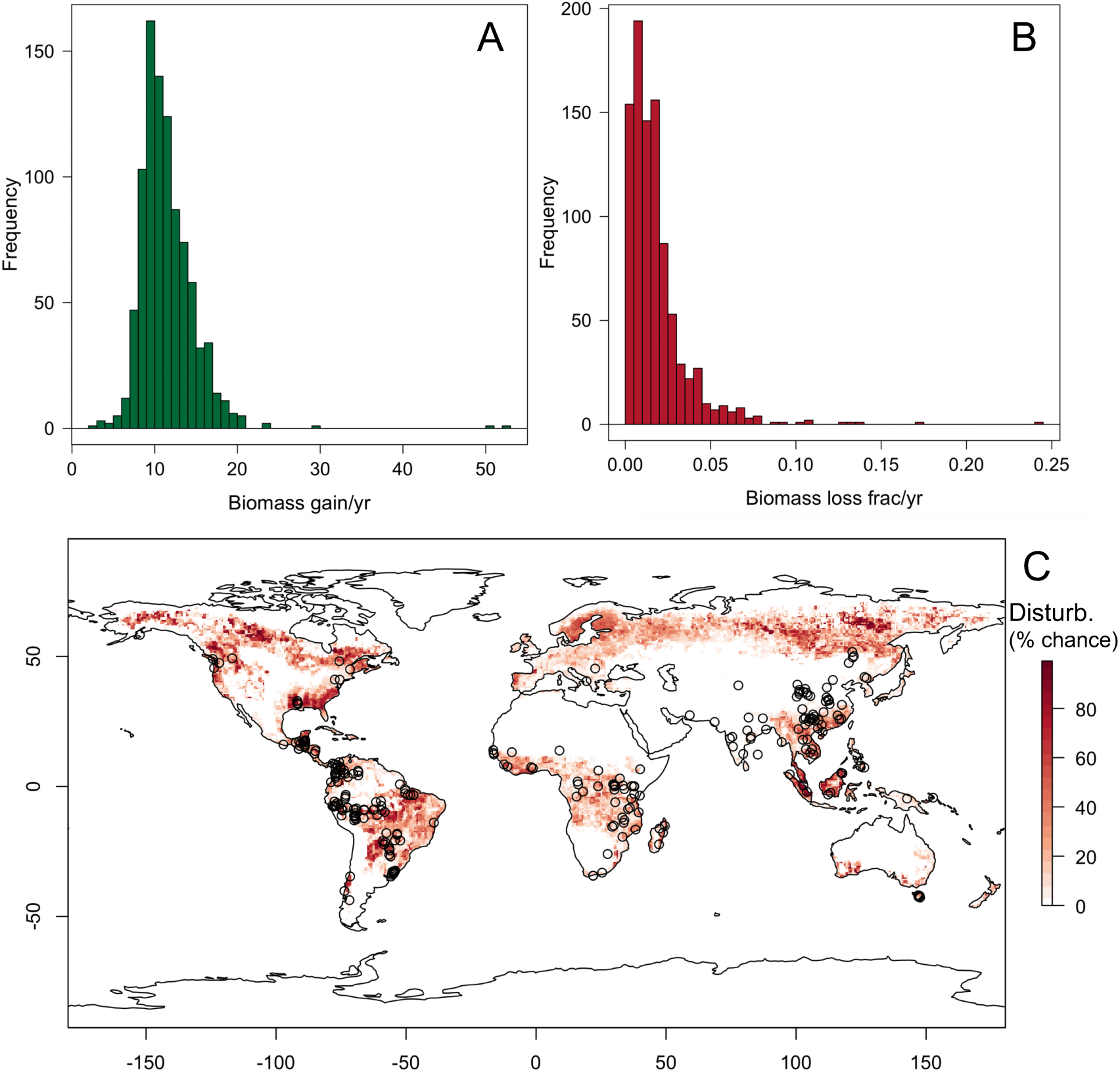
Key inputs of the simple forest model and location of current Verra forest offset projects. (A) Distribution of tropical forest plots’ biomass gain (tons biomass/ha*yr) and (B) biomass loss (fraction biomass/yr) from long-term plot data. These distributions form the basis of the Monte Carlo runs in all analyses. (C) Observed spatial distribution of Verra’s current registered forest offset projects (N=265) overlaid on the 100-yr estimated integrated stand clearing disturbance risk (% chance of at least one stand-clearing disturbance over 100 years) from Anderegg et al. (2022).

**Figure 2:**
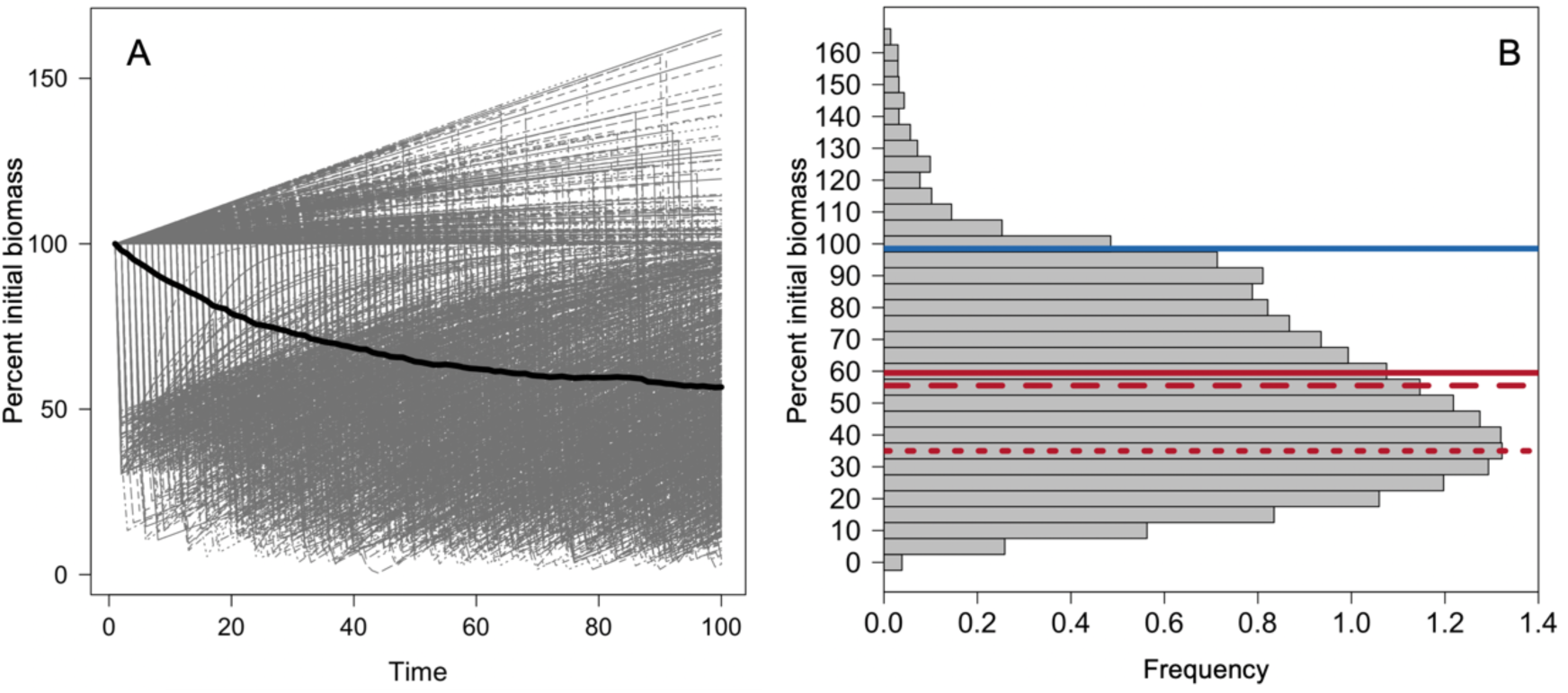
Stand biomass is strongly impacted by moderate and high severity disturbances. (A) Model simulations of the percent initial biomass remaining over a 100 year period in the scenario of biomass losses of 50-70% once every 25-50 years across the 1000 Monte Carlo iterations (black line is the mean). (B) Distribution of the percent initial biomass remaining at 100 years. Blue line indicates the lowest biomass levels insured with the current baseline buffer pool from Verra for this scenario. Solid red line indicates the buffer pool levels needed to insure the mean of the distribution, thick dashes is the median, and thin dashes is the mode.

Considering all disturbance severity and frequency categories, current buffer pool contributions dramatically underestimate the carbon cycle impacts of disturbance, especially at high levels of carbon losses and higher frequencies of disturbance (Fig. 3). For older stand ages (improved forest management and avoided conversion projects), the default Verra buffer pool contribution was inadequate to insure the mean biomass trajectory in 75% of scenarios and inadequate to insure 80% of biomass trajectories (i.e. 80^th^ percentile of disturbance risk, 20^th^ percentile of biomass remaining) in 87.5% of scenarios (Fig. 3). We posit that a buffer pool that does not include the mean project biomass trajectory is substantially undercapitalized and that generally a buffer pool may want to aim to insure >80-90% or more of projects’ biomass trajectories, though this is ultimately a normative decision.

**Figure 3:**
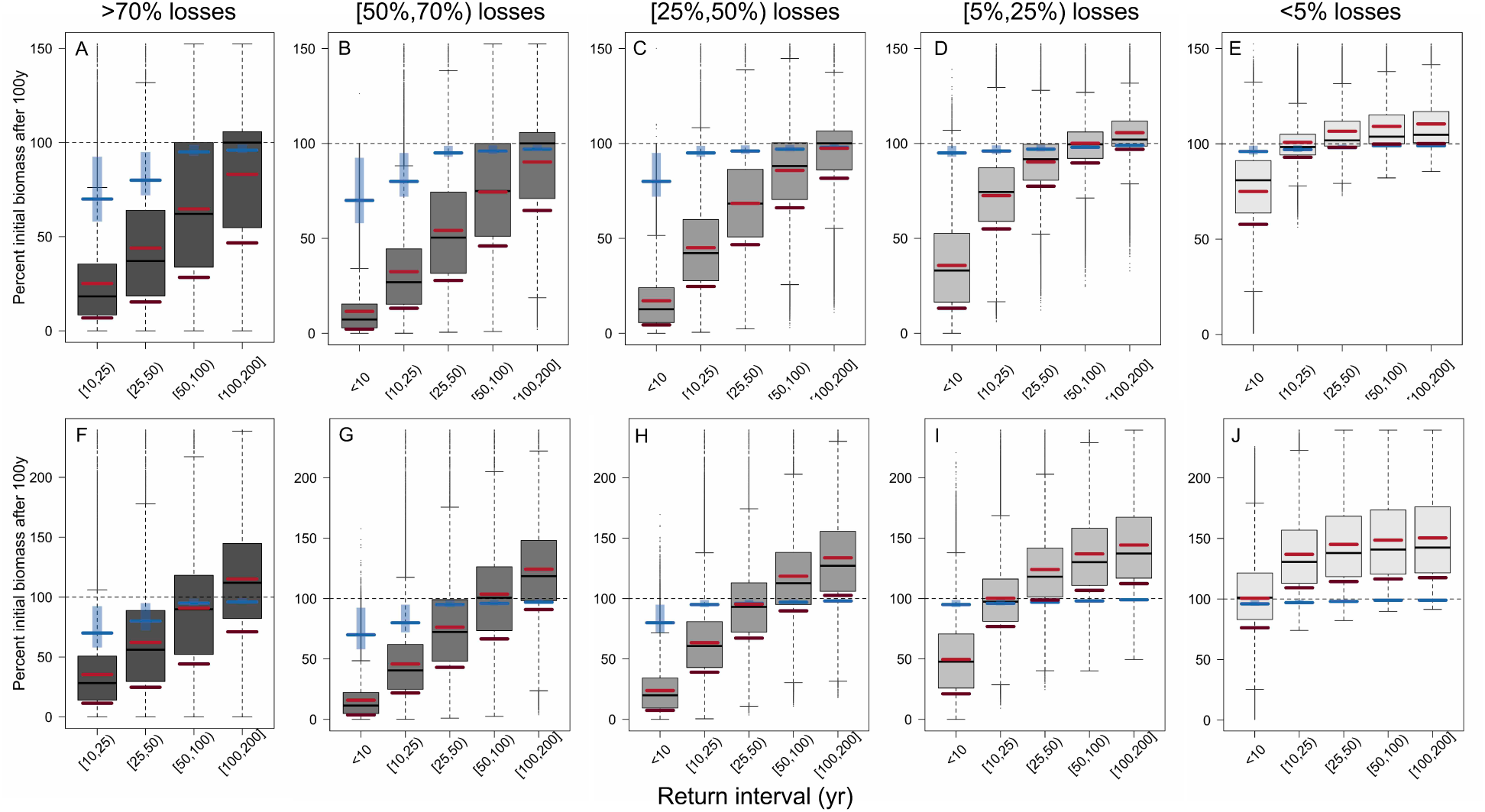
Current buffer pool values are far below levels needed in moderate and high disturbance scenarios. Percent of initial biomass remaining after 100 years as a function of different disturbance severities (darkness of gray from left to right: >70% biomass losses, [50,70%) losses, [25,50)% losses, [5,25)% losses, and <5% losses) and disturbance return intervals (<10 years, [10,25) years, [25,50) years, [50,100) years, and [100,200] years). Top row (A-E) is for average stand age of Improved Forest Management projects (189 years) and bottom row (F-J) is for average stand age of afforestation/reforestation projects (71 years). Blue lines indicate the lowest biomass levels insured with the current baseline buffer pool from Verra and polygons show the range of possible buffer pool contributions. Light red bars indicate the buffer pool levels needed to insure the mean of the distribution in each scenario and dark red bars the 80%ile of the distribution.

Younger forests need modestly lower buffer pools compared with older forests due to faster growth rates (Figs. 3-4). For younger forest ages (e.g. reforestation/afforestation projects), 67% of disturbance scenarios had buffer pool contributions that were inadequate to insure 80% of biomass trajectories (Fig. 3F-J). Further, current buffer pool estimates were still not adequate to insure younger forests (e.g. 50 years old) in moderate and high disturbance regimes (Fig 4B). Of existing Verra projects, there was a bimodal age distribution with reforestation/afforestation projects clustering at low stand ages; improved forest management and avoided conversion projects clustered at high stand ages (Fig 4A). We note that the global forest age product used (Besnard et al., 2021) has a cap at 300 years old and thus does not meaningfully distinguish values above that age.

**Figure 4:**
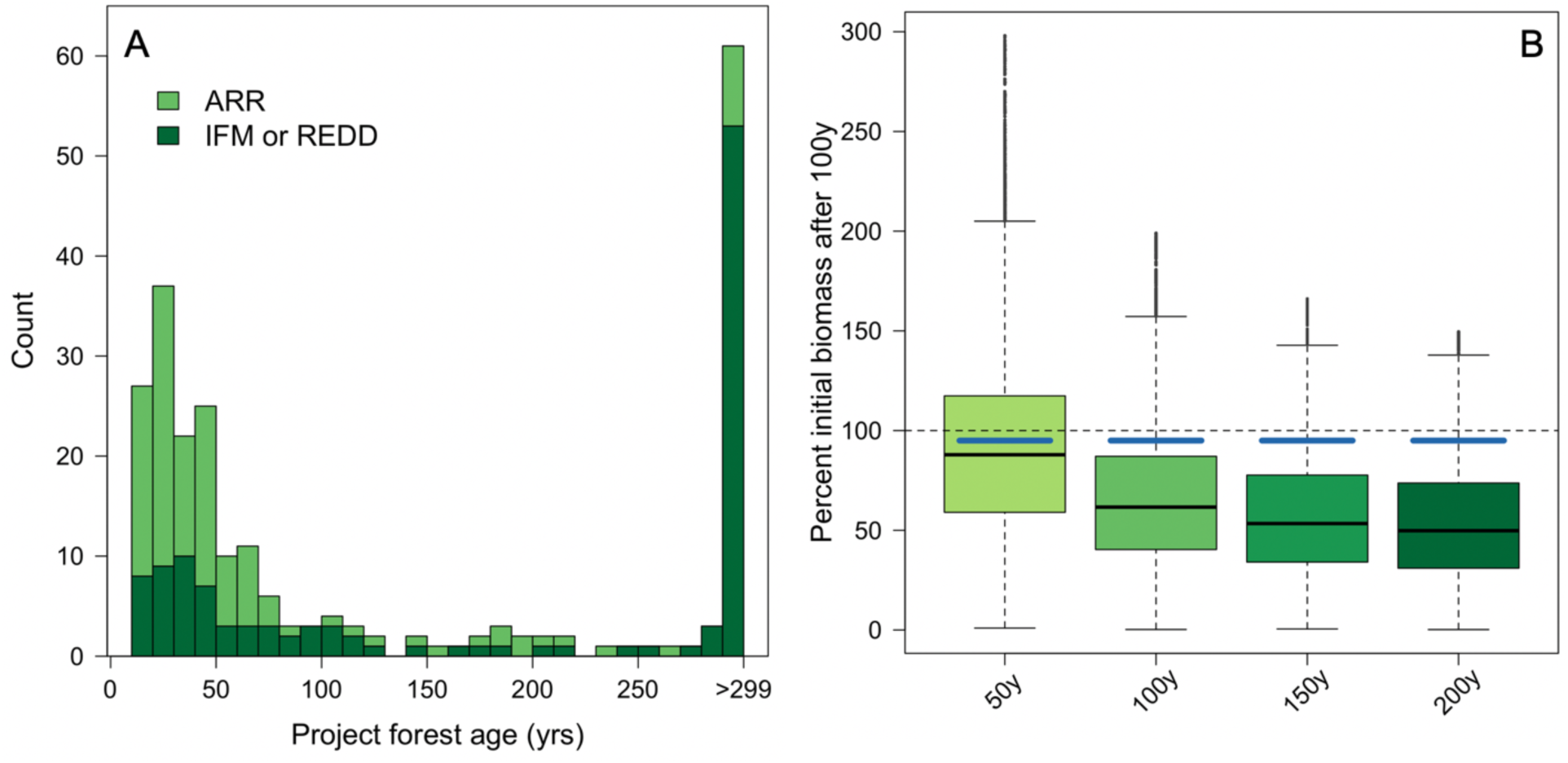
Effects of forest age on simulated biomass trajectories. (A) Forest age distribution for existing registered Verra projects globally for afforestation/reforestation (ARR) and improved forest management or avoided forest conversion in reducing emissions from deforestation and degradation (IFM or REDD). (B) Percent of initial biomass remaining after 100 years as a function of different stand ages (50, 100, 150, and 200 years old) at hypothetical project initiation in the same disturbance scenario as Figure 2 (biomass losses of 50-70% once every 25-50 years). Blue lines indicate the lowest biomass levels insured with the current baseline buffer pool from Verra.

Finally, with the aim of improving science-based policy and protocols, we provide as supplemental tables the buffer pool contributions needed to cover the mean and 80^th^ percentile of forest biomass trajectories based on our analyses in tropical forests (Tables S2, S3) and a global map of stand-clearing disturbance return intervals based on forest losses between 2002-2014 that can be directly incorporated into any updated tool or table as a conservative estimate of combined disturbance risk (Fig S1; see Data statement). We note these estimates exclude land-use change, include forest management activities, and are very likely underestimates of the true risk because they do not incorporate climate change-driven trends in disturbance (see Anderegg et al., 2022 for full details).

## 4. Discussion

We leveraged a forest biomass model and extensive tropical long-term forest plots to probabilistically quantify the impacts of disturbance on 100-year biomass trajectories compared to a widely-used tool in forest offset projects. We find that the buffer pool contributions in Verra’s risk tool are inconsistent with carbon cycle science in the vast majority of disturbance scenarios, especially at moderate or high frequency or severity of disturbance. The inadequacy of the current buffer pool estimates is much larger for older forests, such as those likely to be enrolled in avoided conversion or improved forest management, but still substantial in younger forests. This central finding that current forest offset protocols do not adequately consider risks to forest permanence aligns with substantial recent work in other regions and globally (W. R. Anderegg et al., 2020, 2022; Badgley, Chay, et al., 2022; Coffield et al., 2021; Cooley et al., 2012; Galik & Jackson, 2009; Gren & Aklilu, 2016; Hurteau et al., 2009; Wu et al., 2023).

Moving forward, nature-based climate solutions urgently need to be grounded in rigorous data and carbon cycle science. We provide here clear numbers that would be more likely to insure the mean or most (e.g. 80%) biomass trajectories (Table S2–S3). Crucially, global maps of reversal risk are currently being developed and should be directly incorporated into tools in offset protocols. More detailed research on individual disturbance risks globally, including fire, drought, biotic agents, storms, and sea-level rise, is urgently needed to update such tools (Seidl et al., 2017). Furthermore, factors that either amplify or reduce required buffer pool contributions must both be based on rigorous and region-specific data, which is not the case currently. For example, Verra’s current tool allows 50% reductions in buffer pool contributions if risk “prevention measures are in place” and an additional 50% reduction in buffer pool contributions if “the project has a proven history of effectively containing natural risk” (Verra, 2023). For fire risk, examples of evidence that project developers can provide include fuel removal, establishment of fire breaks, or access to fire-fighting equipment. However, the extent to which these management activities can reduce climate risks, such as fire, is an open scientific question and a 50-75% reduction in fire risk for implementing one or more of these strategies is not based on data. We emphasize that flexibility in allowing project developers to claim buffer pool reductions for poorly-defined management activities (which are additional on top of already undercapitalized buffer pools) is highly likely to lead to an inadequate buffer pool. Furthermore, it’s unclear that risk mitigation for disturbances such as fire or drought or biotic agents is actually effective in many tropical forests (e.g. (Moreau et al., 2022; Nair, 2007)). This highlights that risk reductions in the current protocols lack a solid scientific foundation.

Our analyses present a first-order analyses of the impacts of disturbance on tropical forest carbon dynamics. Our stand biomass model explores a wide range of forest demographic rates but does not include forest composition, mechanistic processes (e.g. competition for light or water, lianas, impacts of severe droughts or pests, effects of rising atmospheric CO_2_ concentrations), or belowground or soil carbon dynamics. More mechanistic and detailed modeling work using dynamic vegetation models and demographic models is needed to explore and further quantify these dynamics and how global change drivers are likely to further influence forest carbon in the 21^st^ century (Brodribb et al., 2020; R. A. Fisher et al., 2018; Walker et al., 2021).

In conclusion, rigorous nature-based climate solutions urgently need 1) to ensure buffer pool contributions reflect carbon cycle science and are based on cutting-edge and publicly-available scientific datasets, 2) to use external and rigorous estimates of disturbance probability and frequency that are standardized by a third party and not chosen by project developers, and 3) to ensure that any risk amplification or deduction factors are based on rigorous science as well. The stand-clearing disturbance return-interval presented here (Fig 1; Supplementary Data) can start to help address #2, but more work that includes climate change trends is urgently needed here. Finally, we note that alternate funding mechanisms for NbCS efforts that are not carbon offsets, such as contribution approaches, are another promising pathway because the uncertainty and rising risks to forest carbon durability are much less of a concern due to the decoupling of forest carbon from fossil fuel emissions (Blanchard et al., In review; Christa M. Anderson et al., 2022). Rigorous science is urgently needed for all NbCS efforts, but the time-horizons of durability required can be shorter for non-offset mechanisms and the consequences of project failure lower for the climate (Blanchard et al., In review; Cullenward et al., 2023).

## Acknowledgements

WRLA acknowledges support from the David and Lucille Packard Foundation, US National Science Foundation grants 1714972, 1802880, 2003017, and 2044937. ATT acknowledges funding from the NSF Grants 2003205 and 2017949 and 2216855, the Gordon and Betty Moore Foundation GBMF11974, and the University of California Laboratory Fees Research Program Award No. LFR-20-652467. GVG acknowledges support from the NOAA Climate and Global Change Postdoctoral Fellowship Program, administered by UCAR’s Cooperative Programs for the Advancement of Earth System Science (CPAESS) under the NOAA Science Collaboration Program award #NA21OAR4310383. CW acknowledges support from the David and Lucille Packard Foundation and the Wilkes Center for Climate Science and Policy of University of Utah.

## Supplementary Tables and Figures

**Table S1:**
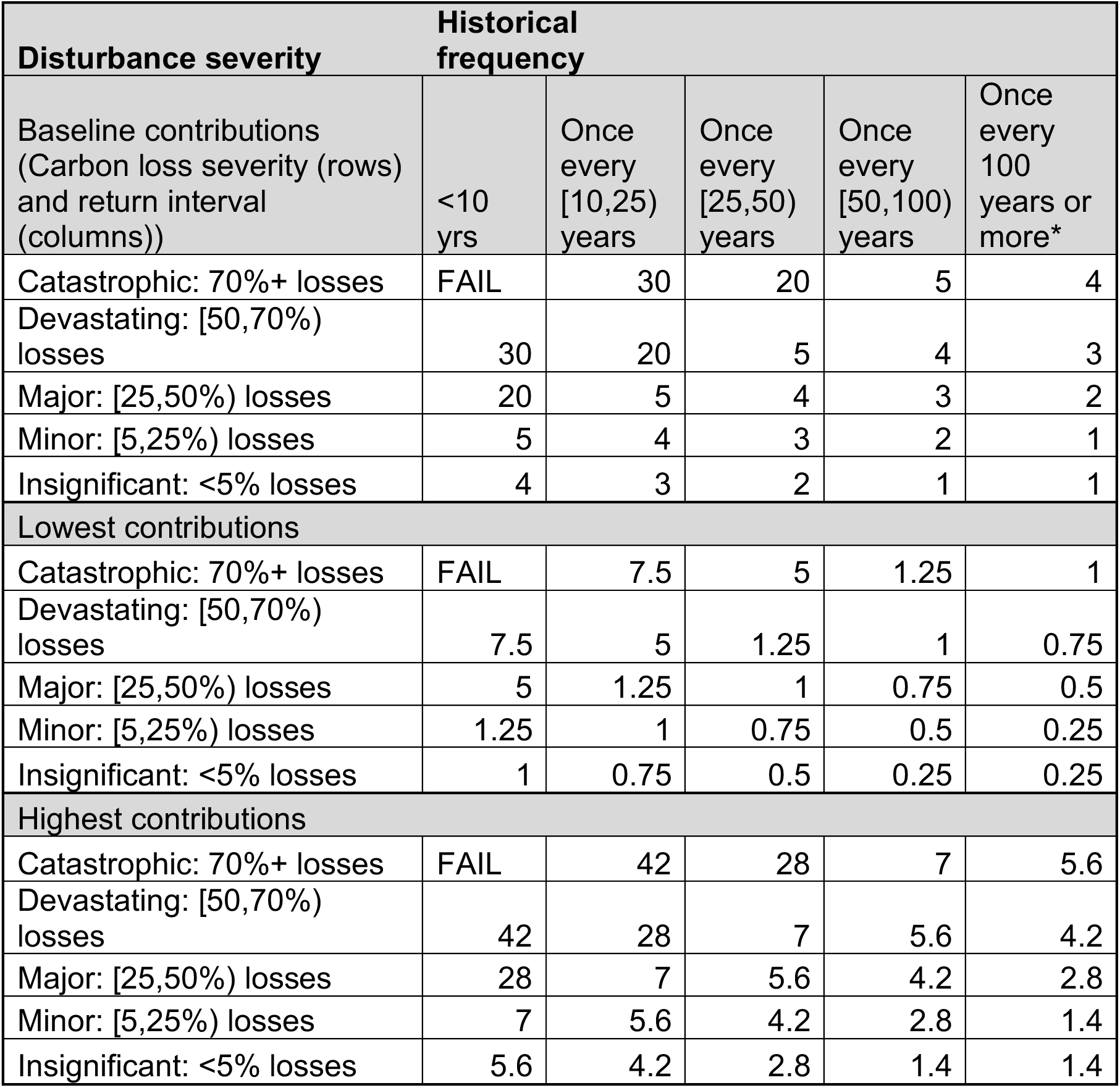
Existing baseline, lowest, and highest buffer pool contributions in Verra’s AFOLU Non-Permanence Risk Tool (reproduced from Table 10 and accompanying equations) (Verra 2023).

**Table S2:**
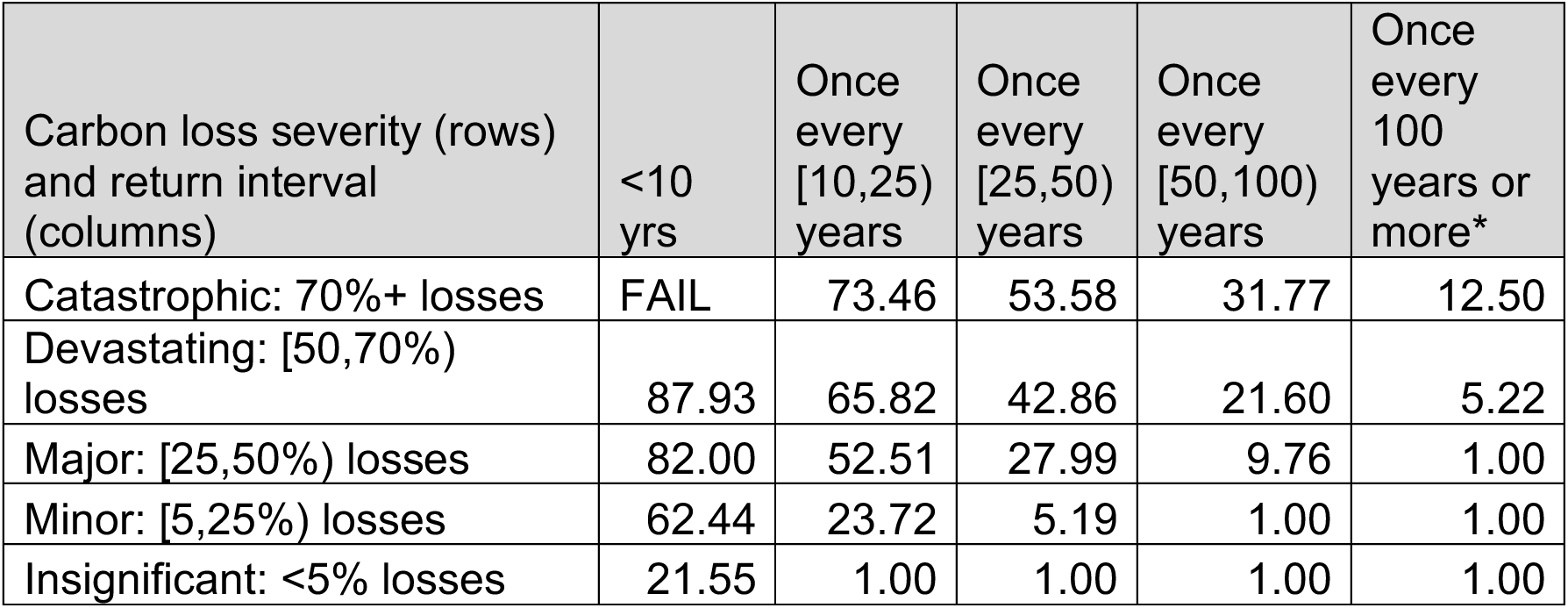
Baseline buffer pool contributions to cover the mean project risk per disturbance type in this analysis.

**Table S3:** Baseline buffer pool contributions to cover 80% of project risk per disturbance type in this analysis.

| Carbon loss severity (rows) and return interval (columns) | <10 yrs | Once every [10,25) years | Once every [25,50) years | Once every [50,100) years | Once every 100 years or more* |
| --- | --- | --- | --- | --- | --- |
| Catastrophic: 70%+ losses | FAIL | 92.42 | 83.19 | 69.17 | 49.41 |
| Devastating: [50,70%) losses | 97.59 | 85.44 | 69.78 | 50.96 | 31.33 |
| Major: [25,50%) losses | 95.09 | 73.10 | 50.16 | 30.36 | 14.38 |
| Minor: [5,25%) losses | 85.59 | 41.54 | 19.00 | 7.07 | 1.00 |
| Insignificant: <5% losses | 39.47 | 4.62 | 1.00 | 1.00 | 1.00 |

**Figure S1:**
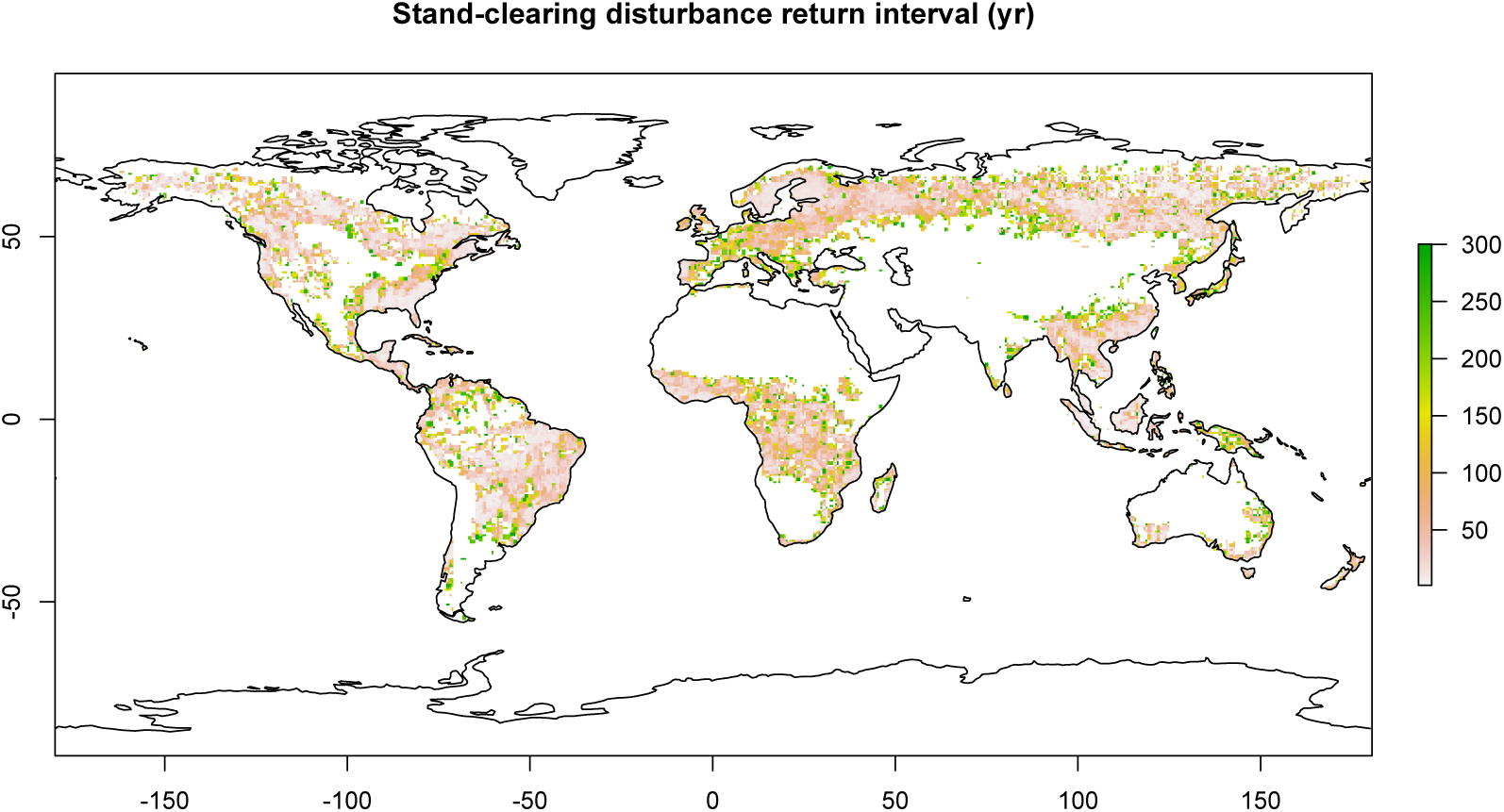
Return interval (yrs) of stand-clearing disturbance estimated from Landsat data from 2002-2014 (full details in Pugh et al., 2019 and Anderegg et al., 2022). Note that stand-clearing disturbance here excludes land-use change but includes all ecological disturbances and management disturbances. Areas with greater than 300 year return-time are displayed as 300 year return-times for visualization purposes. Areas in white are non-forested or a return-time cannot be calculated from the available record due to too infrequent of detected disturbances (e.g. central Amazon) (Pugh et al., 2019).

